# A Standardized Protocol for Sample Preparation for Scanning Electron Microscopy (SEM) to Visualize Extrachromosomal DNA (ecDNA)

**DOI:** 10.1101/2023.08.24.554665

**Authors:** Jillann Madren, Jingting Chen, William Dennis, Christina Ford, Kristen White, Elizabeth Brunk

**Affiliations:** Microscopy Services Laboratory, University of North Carolina at Chapel Hill, Chapel Hill, NC 27516; Department of Biochemistry and Biophysics, University of North Carolina at Chapel Hill, Chapel Hill, NC 27516; Department of Chemistry, University of North Carolina at Chapel Hill, Chapel Hill, NC 27516; Integrative Program for Biological and Genome Sciences (IBGS), University of North Carolina at Chapel Hill, Chapel Hill, NC 27516; Department of Pharmacology, University of North Carolina at Chapel Hill, Chapel Hill, NC 27516; Computational Medicine Program, University of North Carolina at Chapel Hill, Chapel Hill, NC 27516

**Keywords:** Extrachromosomal DNA, double minute chromosomes, scanning electron microscopy, correlative light and electron microscopy, fluorescence microscopy

## Abstract

Extrachromosomal DNA (ecDNA) are large (∼kilo to megabase) acentric, atelomeric, circular DNAs that are established cytogenetic markers for malignancy, hard-to-treat tumors and drug resistance. Often referred to as double minute chromosomes, ecDNA have been studied since the 1960s, primarily through molecular biology techniques, such as cytogenetics and fluorescence microscopy. More recently, next generation sequencing technologies present novel opportunities for identifying ecDNA. However, none of these approaches adequately address the architecture, size and composition of ecDNA within single cells. Developing an approach to systematically visualize ecDNA, confirm their circular architecture and determine their ultrastructure and composition is an urgent, unmet need. This work presents a protocol for visualizing ecDNA at high resolution using scanning electron microscopy (SEM). To this end, we have optimized an end-to-end procedure that involves preparing, processing and visualizing metaphase spread samples. This protocol was tested on five human cancer cell lines (COLO320DM, NCIH716, NCIH2170, SKGT2, SNU16), four of which express ecDNA in various amounts and one amplifies DNA via a homogeneous staining region (HSR). This work presents a standardized approach to preparing samples and visualizing ecDNA using SEM.

**Significance:** Extrachromosomal DNA (ecDNA) are proposed to have unique molecular traits, which include acentric, atelomeric and circular DNA. Standardized, high resolution microscopy approaches are in high demand to better understand structure-function relationships of ecDNA.

## Introduction

Cancer cells use extrachromosomal DNAs to amplify oncogenes[1] and distribute gene dosage in an uneven manner across the cell population, thereby contributing to tumor heterogeneity[2–4] and drug resistance[5]. EcDNA are autonomously replicating DNAs and a large majority are predicted to be circular[3]. Increasing evidence indicates that a circular architecture might be a key molecular trait that increases ecDNA chromatin accessibility and transcriptional capacity over chromosomes[6]. However, methods to systematically assess molecular architecture have not been adequately developed. In order to assess morphologic architecture at the magnification level needed for ecDNA, scanning electron microscopy (SEM) is the ideal method[7]. To use SEM, numerous steps during the preparation of the biological samples must be carefully considered to maintain integrity of ecDNA and chromosomal DNA. Optimization and standardization of these steps is crucial to advance visualization and characterization of ecDNA architecture and composition.

Structure-function relationships of ecDNA are not yet understood. Open questions in the field include: “What percentage of ecDNA are circular?” “Do ecDNA supercoil?” and “Do size differences among ecDNA govern their molecular (three-dimensional) structure?” These questions, while critical, remain challenging to address because we currently lack standardized methods capable of resolving ecDNA structure at high resolution. This is mainly due to the small sizes and molecular makeup of ecDNAs, which push the limits of detection for most traditional and modern molecular technologies. Not only are they small (10Kb - <4Mb), ecDNA are heterogeneous in size, shape and composition. Their DNAs are typically composed of fragments of chromosomal DNA, which can multimerize, leading to a wide range of unique breakpoints where genes from different chromosomes are suddenly next to one another[8,9]. Their sequences contain satellite regions[10] and are otherwise indistinguishable from chromosomal DNA, which limits the capacity of next generation sequencing to determine molecular structure. Further, while fluorescence *in situ* hybridization (FISH) has been a gold standard in the field for characterizing ecDNA, restriction in the magnification potential of fluorescence microscopy limits the capacity to determine ultrastructure.

Several techniques provide the means of direct observation of biological samples at high magnification [11]. The most widely accepted modalities to achieve this type of resolution are transmission electron microscopy (TEM), used to categorize internal structures, and scanning electron microscopy (SEM), used for surface applications. While there have been significant advancements in the instrumentation of SEM, there has been significantly less attention paid to sample preparation. The basic tenets of sample preparation for SEM have remained largely unchanged since its inception in the 1930s. These tenets include appropriate fixation of the sample material, contrast enhancement via heavy metal staining, and dehydration, as well as general SEM principles such as sample conductivity, manipulation of the electron beam optics, and detector type [7,12]. For biological samples, this typically includes chemical fixation in paraformaldehyde and/or glutaraldehyde, the use of osmium tetroxide due to its lipophilic nature making it enhance cellular membranes, and critical point drying to best preserve structure and protect from shrinking. The physical and molecular characteristics of ecDNA make it a unique biological specimen, bringing to question the common practices typically used for SEM of biological samples. In this paper we examine these tenets in detail and compare them to practices used in other forms of electron microscopy that are not standard practice in conventional SEM. We present a comparison of these techniques and present an optimized method for viewing ecDNA via SEM.

## Materials and Methods

### Metaphase Spread Preparation

Condensed chromatin in metaphase is crucial for karyotyping. To accurately profile individual ecDNA and chromosomes, SEM karyotyping requires metaphase spreads. A summary of this stage is shown by Figure 1(A). There are three sections to obtain metaphase spreads: 1) arresting cells at metaphase, 2) hypotonic swelling, and 3) fixation of cells.

**Figure 1.**
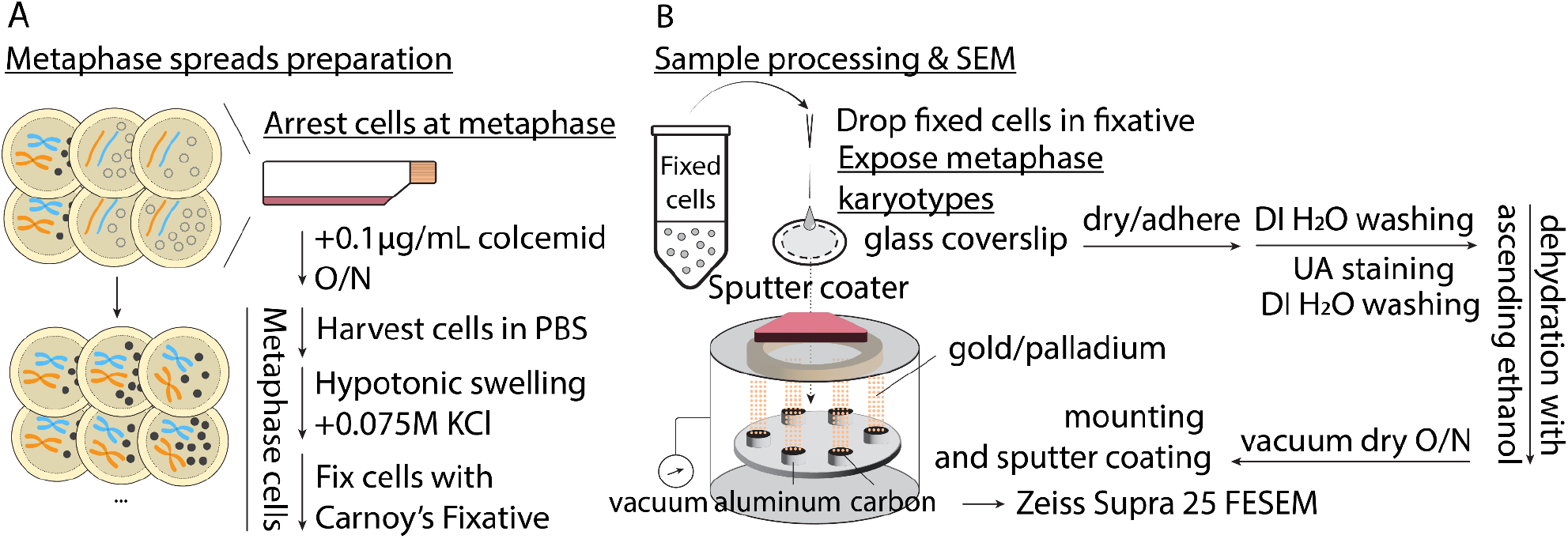
(A) Metaphase chromosome spread preparation. Cultured cells are a mix of non- and mitotic cells. To sync cells at metaphase, cell division is arrested by colcemid, an inhibitor of spindle formation. Cells arrested at metaphase are harvested and washed by PBS, swollen by hypotonic 0.075M KCl solution, and fixed in methanol-acetic acid Carnoy’s fixative. (B) Metaphase samples are prepared for SEM by dropping fixed cells onto a glass coverslip and allowing them to adhere. Cells are stained with uranyl acetate and dehydrated through a series of ascending ethanol solutions, then dried overnight in a vacuum chamber. Coverslips are sputter coated and viewed via FESEM.

First, cells are arrested at metaphase by overnight colcemid treatment (12-20h) at 0.1 μg/mL (10 μg/mL Colcemid Solution, FUJIFILM Irvine Scientific) in cell culture media, when cells are ∼70% confluent. Colcemid prohibits spindle formation, maintenance and cytokinesis, causing metaphase cell arrest. Cells are harvested per standard cell culture procedure with minor alterations. For semi-/adherent cells, metaphase cells are loosely attached and can easily escape into media or PBS wash. To retain cells released in suspension, keep the colcemid-spiked media and PBS wash in a conical tube for quenching trypsinization. Thoroughly resuspend harvested cells in 1 mL of 1x PBS by pipetting. Transfer the suspension to 1.5 mL microcentrifuge tube(s) and each tube should have ≤ 6 million cells. Immediately centrifuge the suspension at 5000 rpm for 2 minutes and aspirate the supernatant.

The second section involves incubating cells with 600 μL of <u>pre-warmed</u> 37°C 0.075M KCl (Gibco). Add it dropwise onto the pellet and gently pipette 3-4 times followed by mild tapping to fully resuspend. Incubate the suspension at 37°C for <u>15 minutes</u> in a water/bead bath. Swollen by osmotic gradients from 0.075M KCl, cells become fragile and their membranes easily rupture. To avoid cells over-/under-swelling and breakage, tightly control incubation time, and no intense handling like pipetting or vortex hereafter.

The third section involves cell fixation. Prepare Carony’s fixative (3:1 methanol:glacial acetic acid) in a fume hood while waiting for incubation. To quench the reaction, directly add 600 μL of <u>freshly prepared</u> fixative dropwise per tube. Immediately centrifuge at 5000 rpm for 2 minutes. To fix cells, a) remove and leave a droplet of the supernatant (100-200 μL). b) Gently tap or flick the tube until the pellet is fully resuspended. c) Add 600 μL of fixative dropwise, resuspend again by flicking, and centrifuge at 5000 rpm for 2 min. Repeat steps a-c twice. Finally, repeat a-b once. Depending on pellet size, add 0-1 mL of fixative dropwise to make a 5-7 million cells/mL suspension. This is the optimal density that ensures discernible spacing for individual cells and chromosomes/ecDNAs therein when dropped on a coverslip.

### SEM sample preparation

A volume of 10 μL of metaphase spread was dropped onto a coverslip (Poly-d Lysine, Neuvitro, 12 mm round or Fisherbrand, 12 mm round, that were cleaned by rinsing three times in 100% ethanol and allowed to dry completely) from a height of at least 50 cm and allowed to adhere for various lengths of time. For contrast enhancement, coverslips were washed three times with deionized water (DI H_2_O) and stained with 1% aqueous osmium tetroxide (OsO_4_) for 30 minutes, 4% uranyl acetate (UA) for 2 minutes, or a combination of the two, with 3x DI H_2_O rinses between and following these steps. For samples that underwent ethanol dehydration, an increasing gradient of ethanol (EtOH) (30%, 50%, 75%, 90%, 100%, 100%, 100% for 10 minutes each) was used before undergoing further dehydration by either critical point drying (CPD), 2x hexamethyldisilazane (HMDS) for 10 minutes followed by air drying, or air drying in a vacuum chamber overnight. All sample preparations are summarized in **Table 1**.

**Table 1.** Parameters of each sample analyzed. OsO4 is osmium tetroxide, UA is uranyl acetate, EtOH is ethanol, HMDS is hexamethyldisilazane, and CPD is critical point drying. A representative image of each sample is listed by its corresponding Figure Number.

| Prep | Figure | Adhere | Stain | Dehydration | Sputter | Thickness |
| --- | --- | --- | --- | --- | --- | --- |
| 1 | 3F | 10 min | OsO<sub>4</sub> | EtOH + CPD | Post | 5 nm |
| 1 | 3E | 10 min | None | EtOH + CPD | Post | 5 nm |
| 1 | 3B | 10 min | OsO<sub>4</sub> | EtOH + Air | Post | 5 nm |
| 1 | 3A | 10 min | None | EtOH + Air | Post | 5 nm |
| 1 | 2B | 5 min | UA | Air | Post | 2 nm |
| 1 | 2C | 5 min | UA | Air | None | None |
| 1 | 2A | 5 min | UA | Air | Pre | 2 nm |
| 2 | 3D | O/N | OsO<sub>4</sub> | EtOH + HMDS | Post | 2 nm |
| 2 | 3C | 10 min | UA | EtOH + HMDS | Post | 2 nm |
| 3 | 4A | 10 min | OsO<sub>4</sub> + UA | Air | Post | 2 nm |
| 3 | 4C | 10 min | OsO<sub>4</sub> + UA | EtOH + Air | Post | 2 nm |
| 3 | 4B | 10 min | UA | Air | Post | 2 nm |
| 3 | 4D | 10 min | UA | EtOH + Air | Post | 2 nm |

### SEM imaging

Coverslips were mounted on 13 mm diameter aluminum stubs with carbon adhesive tabs and sputter coated with a 60:40 gold:palladium alloy using a Cressington 208HR Sputter Coater (Ted Pella Inc, Redding CA). Pre-sputter coated coverslips were made prior to metaphase drop and adherence; post-sputter coated coverslips were made at the conclusion of sample processing. Images were obtained using a Zeiss Supra 25 FESEM operating at 3-5 kV using an InLens or SE2 detector, 20 μm aperture, and approximate working distances of 5 to 10 mm (Carl Zeiss Microscopy, LLC, Peabody, MA).

### Correlative light and electron microscopy (CLEM)

A volume of 10 μL of metaphase spread was dropped onto a gridded glass coverslip or 35 mm coverslip dish of variable materials and grid space sizes (MatTek, Ashland, MA, and Ibidi USA, Inc Fitchburg, Wisconsin) from a height of at least 50 cm and allowed to adhere until dry. Samples were washed three times in deionized water, then incubated with a 1.25 μg/mL solution of DAPI for 2 minutes followed by three rinses in deionized water. For imaging, we used an Andor Dragonfly 200 spinning disc confocal microscope, in widefield mode, equipped with an HC PL APO 100x/1.40 OIL CS2 objective, Zyla Plus 4.2MP sCMOS camera, and controlled by Fusion software version

2.3.0.54. Pixel size was 6.28 um. A 405 nm laser was used to excite the sample and emitted fluorescence was captured in the 424-466 nm range using an Andor Dragonfly Semrock TR-DFLY-F445-046 filter for DAPI. Overview 10x DAPI images were captured and used as a map to select regions of interest in the grids. Images for overlays were captured at 100x magnification in both singular areas or variable sized tiled regions, with a 10% overlap and stitched in high quality mode using Fusion software.

Coverslips were then stained with 4% uranyl acetate for 2 minutes and dehydrated through an increasing series of ethanol (30%, 50%, 75%, 90%, 100%, 100%, 100% for 10 minutes each) before air drying in a vacuum chamber overnight. CLEM coverslips were mounted, sputter coated, and imaged as described above using manual matching of labeled grid squares to image correlated clusters of chromosomes from the fluorescent images.

Overlays were created using the BigWarp[13] plugin for ImageJ[14]. The SEM image was set as the Target image and the fluorescent image was set as the Moving image. We used manual landmarks and transformed using the Affine model.

## Results

Here, we detail the steps required to prepare a biological sample for SEM including fixation, coverslip or substrate selection, contrast enhancement, dehydration, and coating with a metal to improve the electrical conductivity of the sample[15]. A summary of this process is given in Figure 1(B). We evaluated three critical steps: (i) contrast enhancement via heavy metal staining; (ii) drying to preserve structure and protect from shrinking; (iii) sputter coating to increase electrical conductivity. Additionally, we evaluated imaging criteria in two separate steps, which include: (i) SEM settings; (ii) pairing SEM with Correlative Light Electron Microscopy (CLEM)[16]. Lastly, we determined the effect of sample age (the time between the preparation of the metaphase spread and imaging via SEM). The optimal selection for each step is summarized in **Table 2**.

**Table 2.** Summary of Parameters Tested and Optimal Selection for both Chromosomes and ecDNA.

| Parameter | Options | Chromosomes | ecDNA |
| --- | --- | --- | --- |
| Age of Sample | Freshly prepared, older | fresh | fresh |
| Plating to processing | Until dry, overnight | Until dry | Until dry |
| Contrast Enhancement | None, OsO<sub>4</sub>, UA, OsO<sub>4</sub>+UA | UA | UA |
| Drying Method | Air, HMDS, CPD | CPD | Air |
| Dehydration | EtOH, none | EtOH | EtOH |
| Sputter | Pre, Post, none | Post | Post |
| Sputter thickness | 2nm, 5nm | 2nm | 2nm |
| SEM Accelerating Voltage | 3kV, 5kV | either | either |
| SEM Detector | SE2, InLens | SE2 | SE2 |

### Contrast Enhancement

Osmium tetroxide is the most common secondary fixative used in SEM, following formaldehyde/glutaraldehyde treatment. It acts by reacting with unsaturated acyl chains of membranes and lipids as well as to nucleophilic groups. It has mainly been used in SEM to increase contrast in imaging, but it has also been determined to increase the retention of hydrophobic molecules in tissues and cells.

Our initial experiments determined that the use of osmium tetroxide may not be optimal. Borrowing a technique traditionally used for TEM[17] or serial sectioning SEM[18], we implemented staining with uranyl acetate and uranyl acetate in conjunction with osmium tetroxide. We determined that the optimal contrast enhancement was the use of uranyl acetate alone, but the combination of uranyl acetate and osmium tetroxide produced similar, high quality images. However, the use of osmium tetroxide carries additional exposure, handling, and disposal concerns, as well as increases processing time and sample manipulation.

### Dehydration and Drying

SEM samples must be completely dehydrated prior to imaging. This can be problematic for biological samples that naturally have high water content and are delicate and can be easily damaged in the drying process. Changes in surface tension, which occur when transitioning from the liquid to the gaseous state, can cause damage to the sample and can be prevented through appropriate drying techniques to preserve surface structure. For most biological samples this involves critical point drying, a method that utilizes temperature and pressure to achieve a state in which the liquid and gaseous states of the solvent are indistinguishable and can therefore cross membranes with no surface tension effects.

To determine which drying method was most suitable, we tested three main approaches: (i) hexamethyldisilazane (HMDS); (ii) critical point drying (CPD); and (iii) air drying. We performed each of these techniques after an increasing gradient ethanol dehydration. While HMDS is a chemical drying method reported to mimic CPD, we determined that HMDS was not suitable for this application as it appears the chromosomes expanded to bursting and created a large field of debris on the samples. Critical point drying (CPD) had high structural preservation of chromosomes, but ecDNA were much fewer in number than expected for the tested cell lines. Air drying showed slightly less surface detail on the chromosomes, but retained more ecDNA, which we concluded was an acceptable compromise as the ecDNAs were our area of interest.

Lastly, we tested an air-drying method to determine if ethanol dehydration alone would improve chromosome quality. We found that including an ethanol dehydration process prior to vacuum air drying provided better structural preservation than air drying alone. Chromosomes dried in the vacuum chamber alone with no ethanol dehydration showed artifactual signs of shrinkage and surface collapse.

To compare these drying methods, we measured the diameter of several chromosome pairs for each method tested and compared them to the accepted published measurement of 1400 nm[19]. The method closest to the known value of 1400 nm was uranyl acetate dehydrated with an ethanol gradient then air-dried, which gave an average measurement of 1566 nm.

### Sputter Coating

For SEM, sputter coaters are used to generate a thin film that is electrically conductive and representative of the biological specimen. The main objective of using sputter coaters are to prevent electrical charging, thermal damage and improve secondary electron emission.

Considering that ecDNAs are flat and that DNA is slightly conductive, the need for sputter coating was evaluated. We generated three conditions to optimize sputter coating application: (i) pre-processing, (ii) the conventional post-processing, as well as (iii) no coating. We applied a 2 nm 60:40 Au:Pd to the coverslip for the pre-processing and the conventional post-processing. For the condition without sputter coating, we imaged it directly with only the UA staining. The uncoated coverslip exhibited a high amount of charging and was essentially unviewable. We determined that we could pre-sputter our coverslips, but ultimately decided that a thinner 2 nm coat performed post-processing was the more conservative method, especially for samples that undergo critical point drying (discussed below, data not shown). Other untested alternatives are conductive substrates such as carbon coated or silicon wafers or aluminum or indium tin oxide coated coverglass.

### SEM Microscope Settings

To determine our preferred microscope settings, we imaged the same field of view with SE2 and InLens detectors at 3 kV and 5 kV accelerating voltages. We determined that the best quality image for chromosomes and ecDNA was obtained using the SE2 detector at 3 kV, however 5kV also produced satisfactory images. Additionally, the InLens detector could be useful for finding areas of interest due to the “glow” effect observed. However, the InLens detector resulted in obscured chromosomes and therefore the SE2 detector was the better choice for final imaging.

### Effects of Sample Age

Sample freshness is key for ultrastructural analysis, where samples begin to degrade within 15 minutes if not placed in fixative and that repeated freeze/thaw cycles are detrimental to sample integrity. To test the effects of sample age, we imaged samples that had been stored per protocol at -20°C for several months and samples that were freshly prepared immediately prior to SEM processing and imaging. We determined that the freshly prepared samples retained their chromosome structures better, as well as surprisingly contained more ecDNA molecules. Other untested considerations would be the effects of cell line passaging.

### Pairing SEM with CLEM

Using SEM, ultrastructural analysis is on a gray scale, making it challenging to differentiate ecDNA from random debris. To determine whether we were visually observing ecDNA, we paired SEM with Correlative Light Electron Microscopy (CLEM). CLEM is a fluorescent microscopy technique that allows for higher resolution than standard fluorescent microscopy techniques. Samples can be stained with fluorescent dyes and visualized in conjunction with scanning electron microscopy. Using this technique, we were able to confirm that the small particles we observed were ecDNA composed of nucleic acids and aligned with the expected size and structure of ecDNA.

For our CLEM experiments, we used two different brands of gridded coverslips, in both coverdish and coverslip format, glass and polymeric material, and grid sizes of 50 or 500 μm to determine which product best met our needs. While we liked the labeling for every grid square provided by the MatTek coverslips, we found them more difficult to determine positioning by SEM than with the Ibidi coverslips. We had a strong preference for the 50 μm grid size as it was much easier to correlate in a smaller frame size. We opted for the coverslip over the 35 mm dish because it was the more economical option and the dishes are not necessary for this protocol, although stabilization during the light microscopy needs to be considered. We did not have a strong preference for coverslip material, but the 50 μm coverslips only come in glass in the coverslip format. These options are summarized in **Table 3**.

**Table 3.**
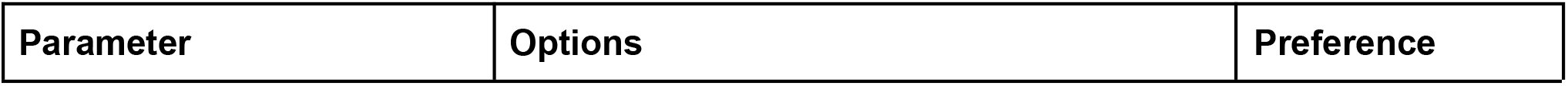
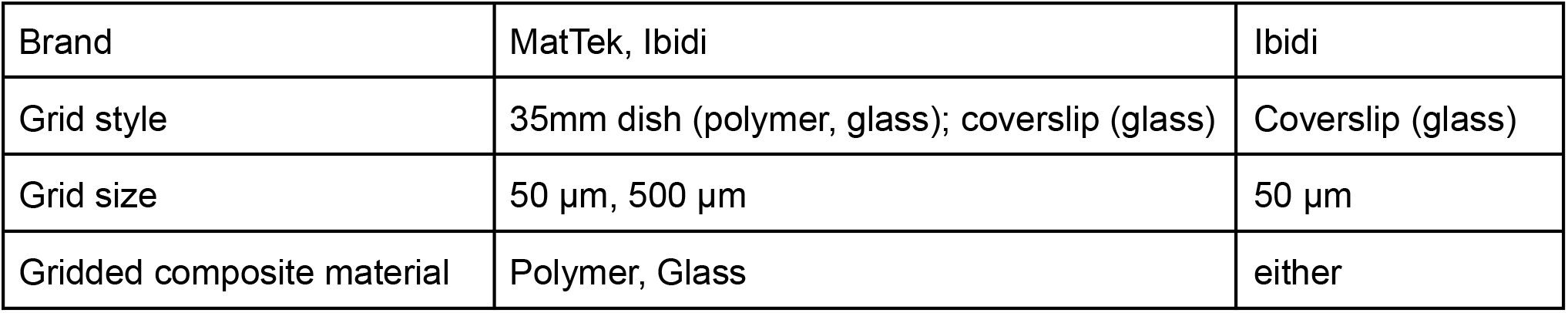
Summary of Gridded Materials Tested and Optimal Selection for Correlative Light and Electron Microscopy (CLEM).

| Parameter | Options | Preference |
| --- | --- | --- |
| Brand | MatTek, Ibidi | Ibidi |
| Grid style | 35mm dish (polymer, glass); coverslip (glass) | Coverslip (glass) |
| Grid size | 50 μm, 500 μm | 50 μm |
| Gridded composite material | Polymer, Glass | either |

## Discussion

Despite the surge of cutting-edge genome technologies, structure-function relationships of ecDNA remain a mystery. For example, does the three dimensional shape and architecture of ecDNA determine its functional role in cancer and cell fitness? Recent studies suggest that many ecDNAs are circular, and that their circularity may play a role in determining how accessible the chromatin is for genes on ecDNA[6]. However, a systematic ultrastructure analysis of ecDNA across cells and cell lines has not yet been carried out. One reason for this is potentially due to the fact that ecDNAs present unique challenges to both conventional and state-of-the-art techniques. Currently, the only approach capable of accurately resolving the higher structure of ecDNA are high resolution microscopy techniques, such as SEM and TEM.

While SEM is considered a standard approach for ultrastructural analysis of biological molecules, the process of sample preparation presents numerous challenges for ecDNA. The physical and molecular characteristics of ecDNA make it a unique biological specimen for SEM. For example, ecDNA are small DNA fragments, which range in size between tens of kilobases to megabases. Therefore ecDNA can “get lost” or mistaken for cellular debris when using SEM. The ideal SEM image would be able to resolve chromosomes, detect ecDNA, be “clean” or free of dust/debris, and retain biological details, such as the ability to view chromosomes in close proximity to ecDNA and separate different cell nuclei. These needs bring to question the common practices typically used for SEM of biological samples and whether they require further optimization to improve the accuracy and visualization of ecDNA structure. In this paper we examined various steps in the preparation of samples for SEM and compared them to practices used in other forms of electron microscopy that are not standard practice in conventional SEM. This work presents a comparison of these different practices to create an optimized protocol for viewing ecDNA via SEM.

An optimized protocol for the ultrastructural analysis of ecDNA will provide new inroads not only for studying structure-function relationships of ecDNA, but will provide a standard method for cores, facilities and other microscopy users that are interested in optimal imaging of ecDNA. Electron microscopes have very special technological demands that prevent most scientific labs from being able to house them. Therefore, this work will be of great value to the broader scientific community, providing knowledge that can be applied across scientific labs through microscopy cores. While numerous conditions were evaluated herein, several other factors may also need to be considered when performing SEM to evaluate ecDNA. These factors include the effect of passage number/cellular age of the sample, and other parameters during cell culture (e.g. the use of trypsin or EDTA). We expect this research to contribute to the ever-increasing number of publications on ecDNA, especially to provide a starting-point for researchers interested in defining structure-function relationships and ultrastructural analyses of ecDNA.

**Figure 2.**
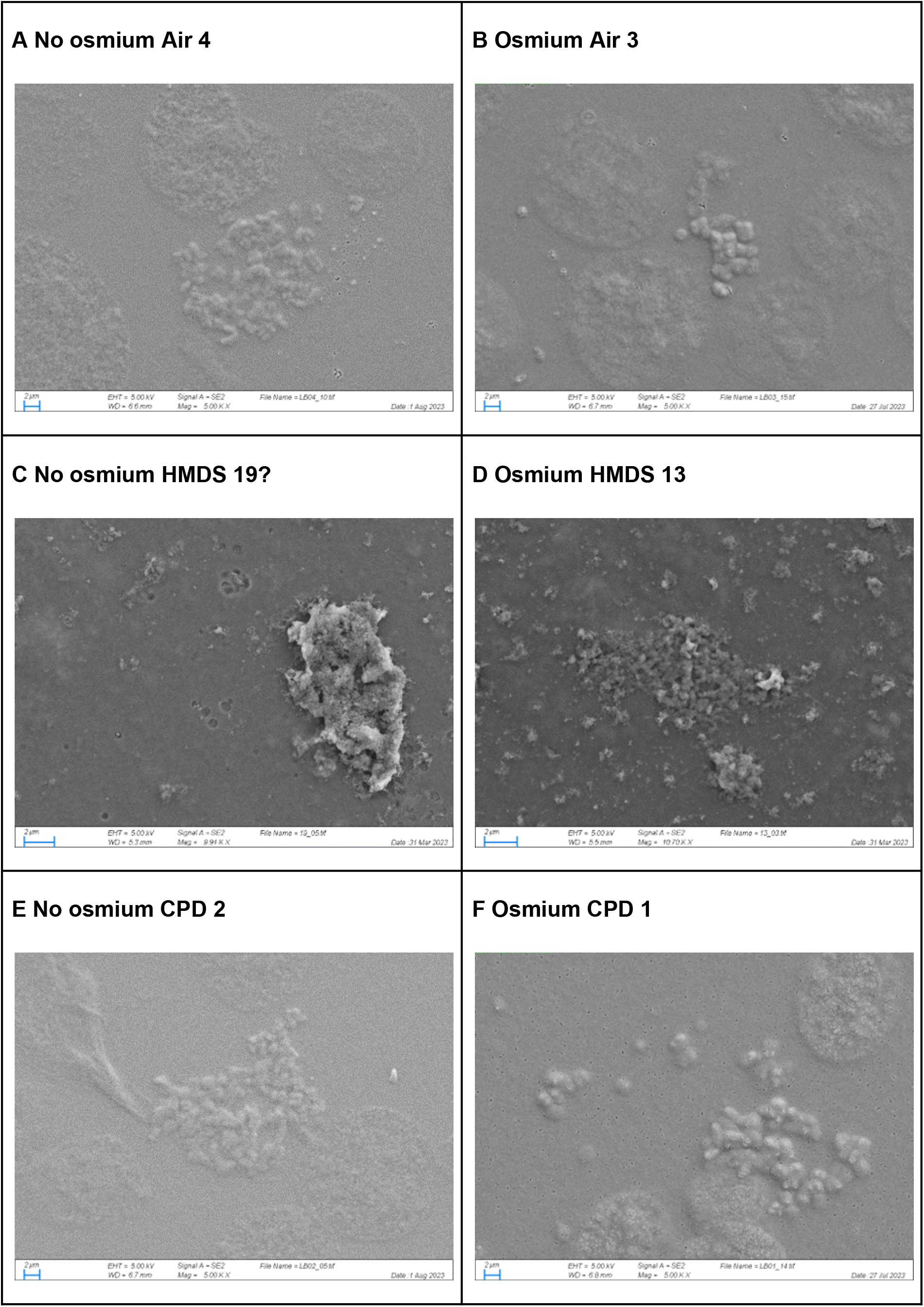
Comparison of chromosomes and eccDNA using conventional SEM protocols. Samples were compared in relation to osmication (B,D,F) or no osmication (A,C,E) and drying methods of vacuum air-dried (A,B), dried chemically with HMDS (C,D), or CPD (E,F). Scale bars are 2 μm.

**Figure 3.**
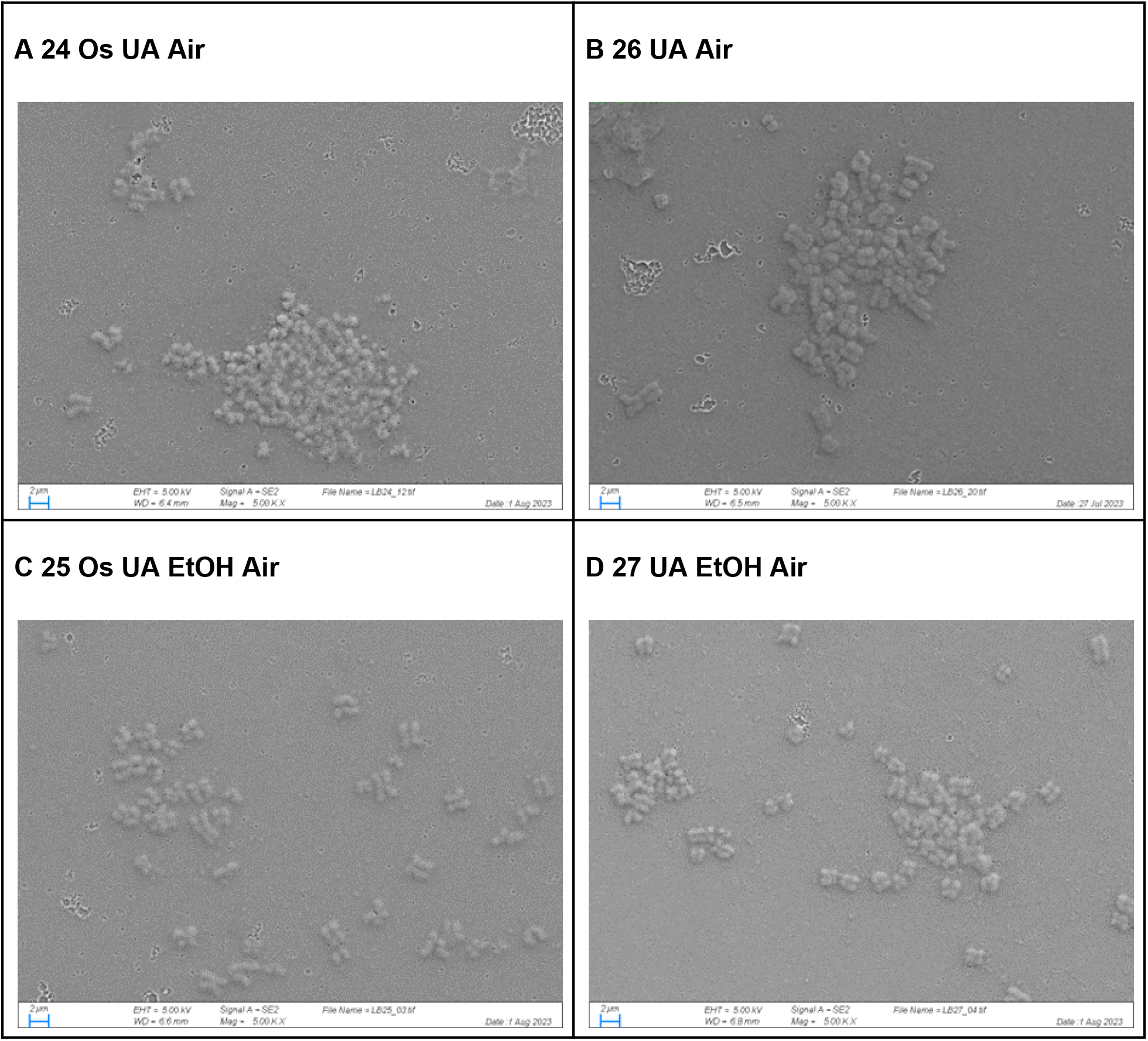
Comparison of contrast agent and dehydration.

**Figure 4.**
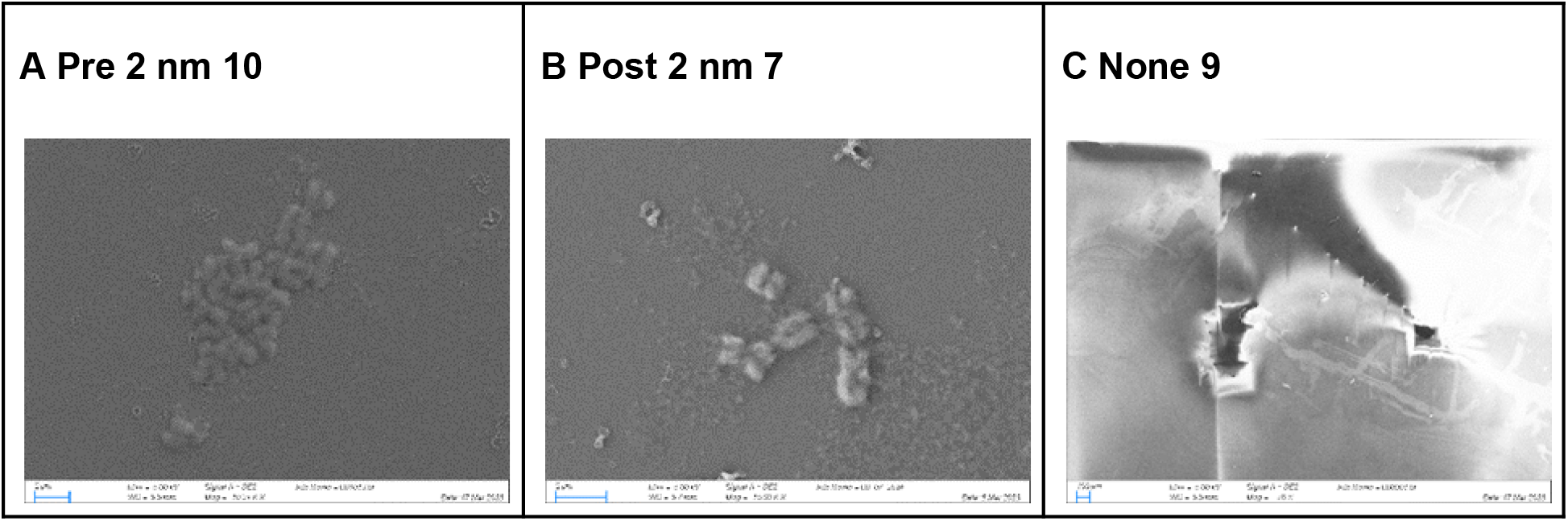
Comparison of sputter coating techniques. Coverslips were coated with 2 nm 60:40 Au:Pd either before (A) or after (B) sample processing or not at all (C). All samples were stained with 4% uranyl acetate for 2 minutes then air dried.

**Figure 5.**
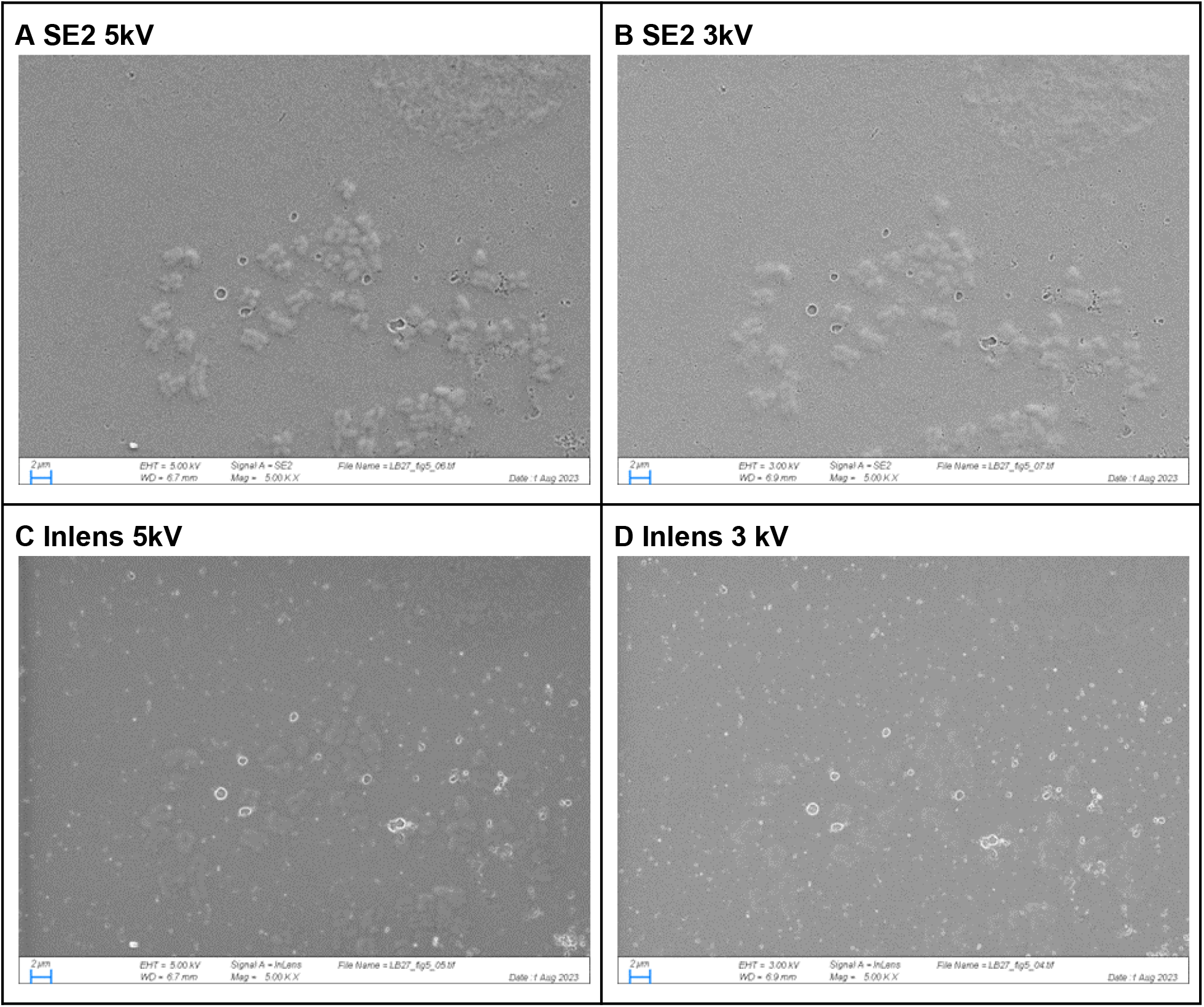
Comparison of SEM detector and accelerating voltage using SE2 detector (A and B) or InLens detector (C and D) and accelerating voltages of 5 kV (A and C) or 3 kV (B and D) using the same field of view with only slight focus modifications. Sample used for these images was processed using the optimized processing protocol of staining with 4% uranyl acetate, dehydrating through an increasing ethanol series, and air drying.

**Figure 6.**
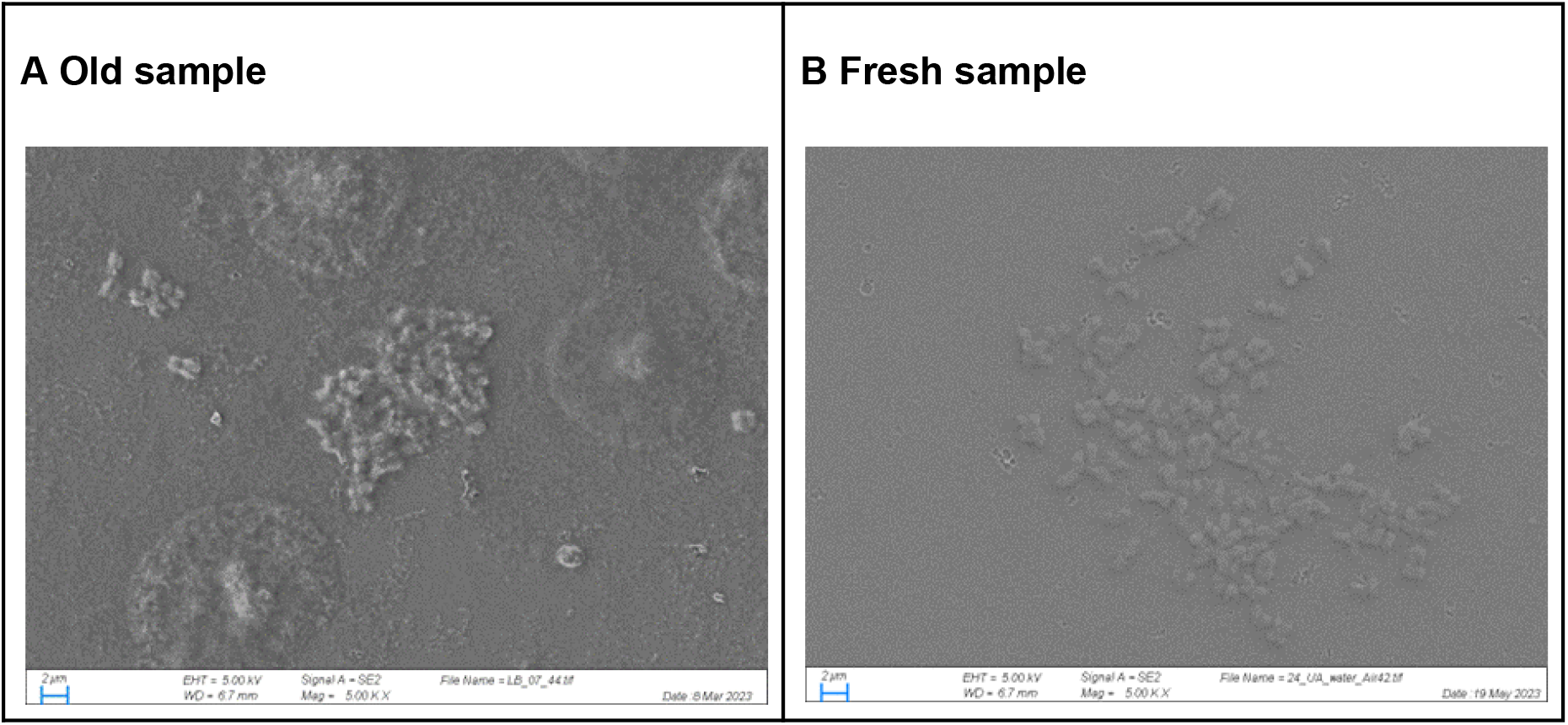
Effect of Sample Age. The optimized processing protocol of staining with 4% uranyl acetate, dehydrating through an increasing ethanol series, and air drying was performed on cells that had been stored per protocol in methanol:acetic acid fixative at -80*C for four months (A) and cells that were freshly prepared the same day as processing (B).

**Figure 7.**
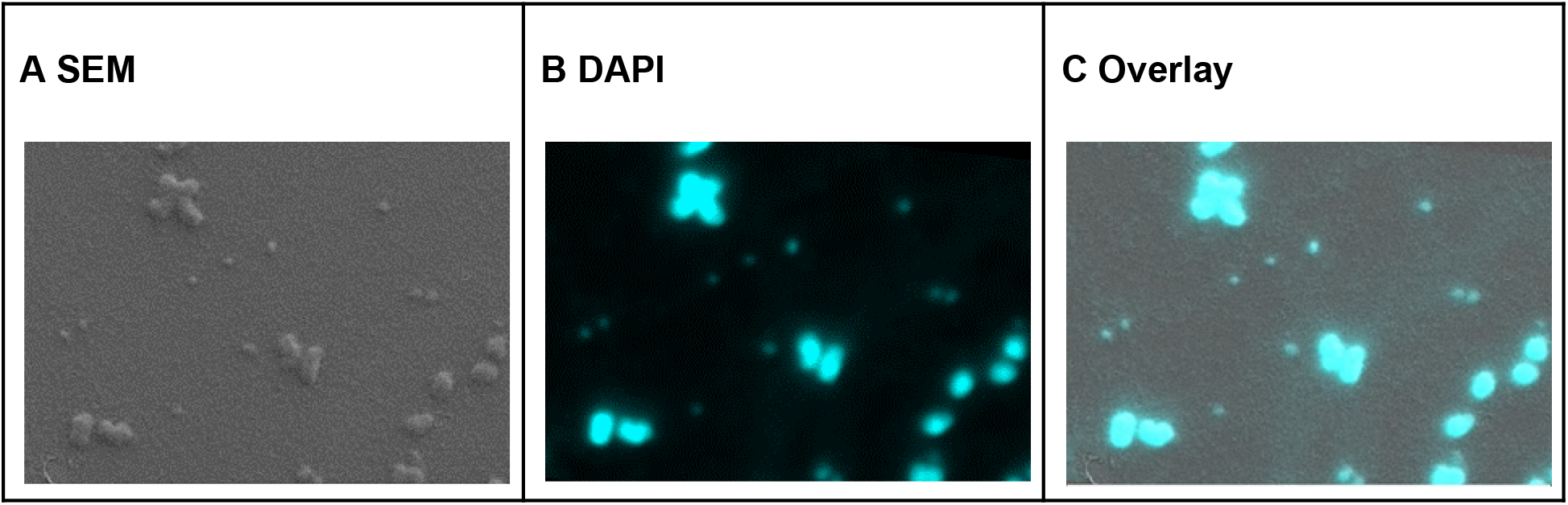
(A) Scanning Electron Microscopy image of chromosomes and ecDNA in the NCIH-716 cell line. (B) Fluorescence microscopy image of the same chromosomes and ecDNA as in (A), nucleic material is stained with DAPI. (C) An overlay of (A) and (B), confirming that the small fragments of ecDNA identified by SEM are DNA.

## Acknowledgements

The authors acknowledge Pablo Ariel for the helpful discussions and input on CLEM.

## Author Contributions

Conceptualization, E.B., K.W., J.M.; methodology, E.B., K.W., J.C., W.D., J.M.; formal Analysis, J.M., K.W., J.C., and E.B.; funding Acquisition, E.B.; investigation, E.B. and K.W.; resources, E.B.; supervision, E.B.; validation, J.M., J.C., E.B., K.W..; visualization, J.M., K.W.; writing—original draft, J.M, E.B.; writing—review and editing, All authors. All authors have read and agreed to the published version of the manuscript.

